# Effect of membrane cholesterol on dynamical properties of solvent molecules

**DOI:** 10.1101/2021.02.19.432063

**Authors:** Hadi Rahmaninejad, Darin D. Vaughan

## Abstract

Membrane lipid composition is a critical feature of cell function, where cholesterol is a major lipid sterol component that influences the membranes physical and electrical properties. The effects of cholesterol on transport properties between adjacent to the cells, especially in junctions formed between cells is not completely understood. These junctions where substances transport and signaling is critical may be affected by modifying the cholesterol composition of the membrane in these junctional regions. Here we show how the cholesterol content in a membrane can regulate these phenomena by changing their effect on transport into and through regions between cell membranes in close proximity. Through geometric and electrostatic effects interaction with substrates, the properties of the fluid between membranes are shown to potentially enforce concentration gradients of dissolved compounds that may be biologically significant.

## Introduction

Biological membranes are composed of a mosaic of proteins and lipids in a fluid-like matrix, with lipids forming a micelle bilayer with proteins embedded or attached to the surface. The precise composition is variable and can range from 20% protein and 80% lipid, to 80% protein and 20% lipid, depending on the cell and tissue. Proteins are generally divided into two classes; integral (or intrinsic) that pass through the membrane entirely, and peripheral (or extrinsic) proteins found on the outer or inner leaflet of a membrane. The membrane-bound proteins are believed to be responsible for varieties of functions, including cell signaling and interacting with ligands [1]. The possible classes of lipids are of a much greater variety, and the relative composition of each class also depends on the particular organellar membrane, as seen in Figure 1.

**Figure 1:**
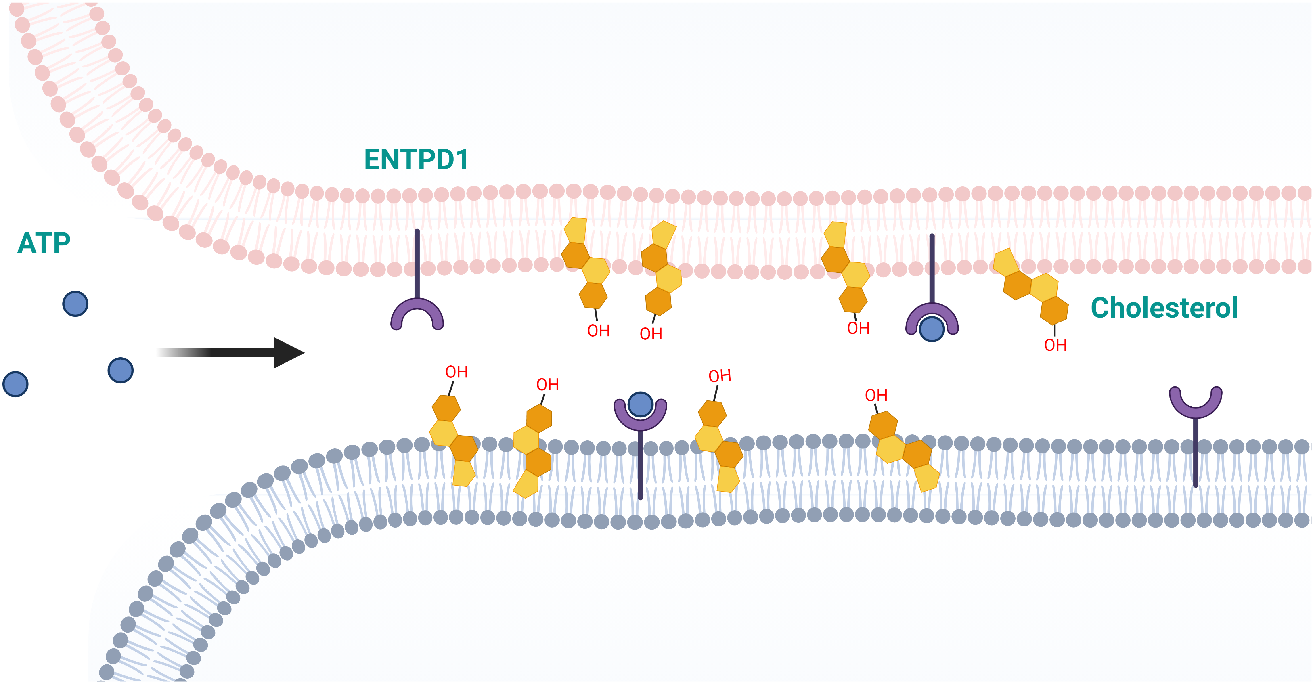
General system design

The geometric properties of the extracellular junctions widely affect the gradient in species concentration by affecting accessible areas. Furthermore, electrostatic interaction can directly affect substances from the external hydrophilic layer of the membrane. It can regulate it through interaction with water molecules as the primary solvent around cells. Hence, phospholipids, as the main constitutes of the membrane, play an essential role, mainly by affecting their physical properties[2]. Cholesterol, as another key component of membrane, not only regulates mechanical and physical properties of membranes, such as their fluidity but also modulates the viscoelastic properties of the membrane which is essential for pathogenesis and cellular organization [3]. It has been shown that cholesterol can increase membrane thickness and area per lipid. They also affect electrostatic interactions by modifying the surface charge density of them [4, 5]. In particular, higher cholesterol content can decrease the average surface charge density of phospholipid membrane [6]. Cholesterol can alter the surface charge density of membranes as high as 25% [7], and consequently modify electrostatic interactions.

Here, we elucidate how cholesterol content modulates the dynamical properties of the surrounding solvent molecules, which in turn influences the transport of solvated substrates. To demonstrate that, we applied molecular dynamic simulations of DMPC/cholesterol bilayers with various cholesterol content to investigate how cholesterol rates the thickness of the membrane and its surface charge density. Then, we computed the diffusion coefficient relatively close and far from the membrane. Therefore, it can be shown that how the accumulation of the change in the diffusion coefficient can result in the variation to the material transport in physiological environments, especially in extracellular junctions, which is critically essential for metabolism.

## Result

To investigate the effects of cholesterol content on the solvent molecules’ transport properties, we must estimate how the rate of cholesterol per area influences the average diffusion coefficient. We performed a series of molecular dynamics simulations and compared cholesterol content effect on geometrical and electrostatic properties of the lipid membrane. Then, we investigated the average diffusion coefficient of solvent molecules near and far from the membrane for different percentages of cholesterol.

### Effective thickness

First, we calculated the z-density of water molecules (see Fig. 2) to estimate membrane boundary based on the density of the water molecules to compute the thickness of the membrane. Here, the x-axis shows the z coordinate, which is perpendicular to the membrane’s surface (see also Fig. 5), and the y-axis represent the z-density of water molecules, which is the total number of water molecules per length of the z-axis in a plane parallel to the x-y surface. Here, blue and red lines represent water molecules’ z-density for 10% and 40% cholesterol contents.

**Figure 2:**
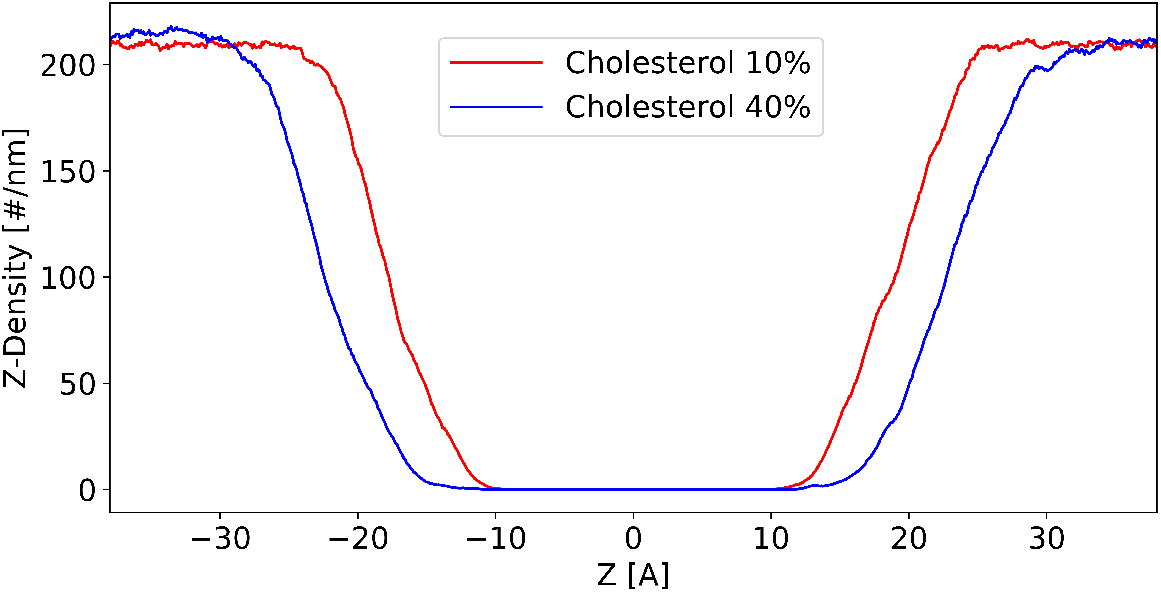
Z-density of Water molecules along the direction normal to the membrane surface (z-axis)

Here, we define an effective thickness for lipid membranes as the total thickness in which there is no net number of water molecules in the z-density plot. Hence, as can be seen, the effective thickness of the lipid membrane increases with the cholesterol content. Other computational and experimental works indicate such proportional relation between membrane thickness and Cholesterol content [8, 9], with computing peak-to-peak distances using electron density profile.

### Surface charge density

We next quantified the average partial charge normal to the membrane surface (along the z-axis in Fig. 3.) Averagely, a more negative charge at the membrane is observed for the membrane with relatively less amount of cholesterol To better show this, we offer a residue view vs. charged view of DMPC membrane, including cholesterol in Fig. 3. The less sharp color of the area with cholesterol indicates how it smoothens the hydrophobicity of the membrane’s surface.

**Figure 3:**
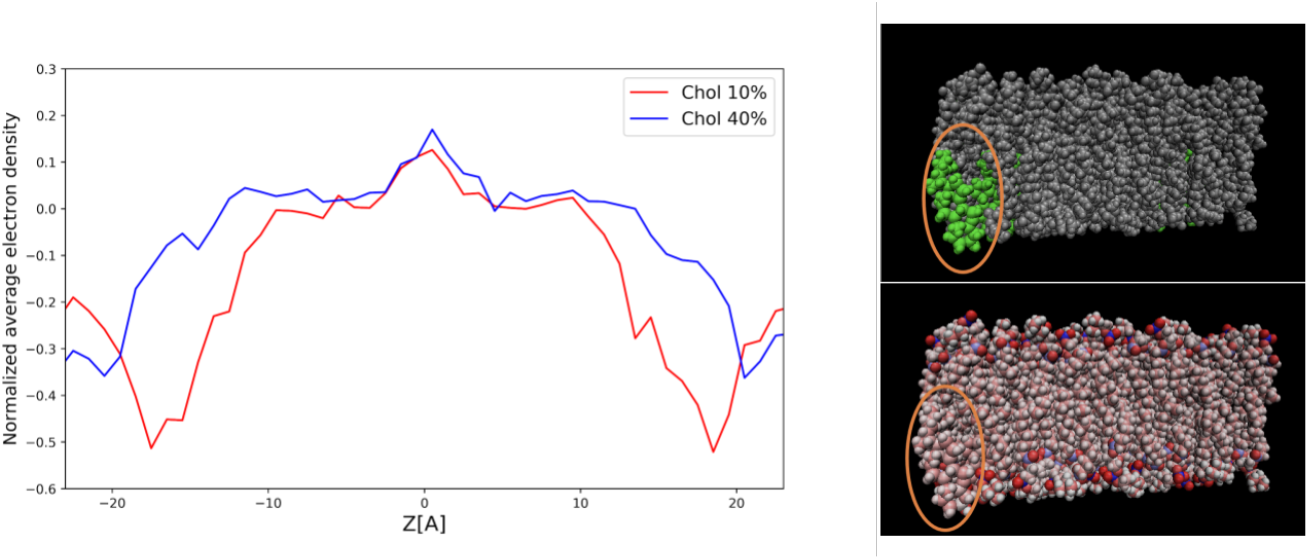
Partial charge of membrane molecules along the z-axis. The top right shows a residue view of the membrane surface with Cholesterol in green, while the bottom right shows the same surface in charge view.

### Diffusion coefficient

We so far demonstrated how cholesterol content alters the effective thickness and average electrostatic charge density of the membrane surface. To investigate how the change in the charge surface density resulting from cholesterol content variation on the dynamical properties of the water molecules around the surface, we computed average diffusion coefficients of water molecules using the Mean Squared Deviation (MSD), both close and far from the membrane. We did this procedure for both cases of high- and a low percentage of cholesterol molecules in Fig. 4. Far from the membrane, the diffusion coefficient is almost independent of the cholesterol content in the membrane, approximately equal to the diffusion coefficient in the bulk case. However, close to the membrane, diffusion is considerably affected by the cholesterol content. More exactly, a higher cholesterol percentage increases diffusivity adjacent to the membrane.

**Figure 4:**
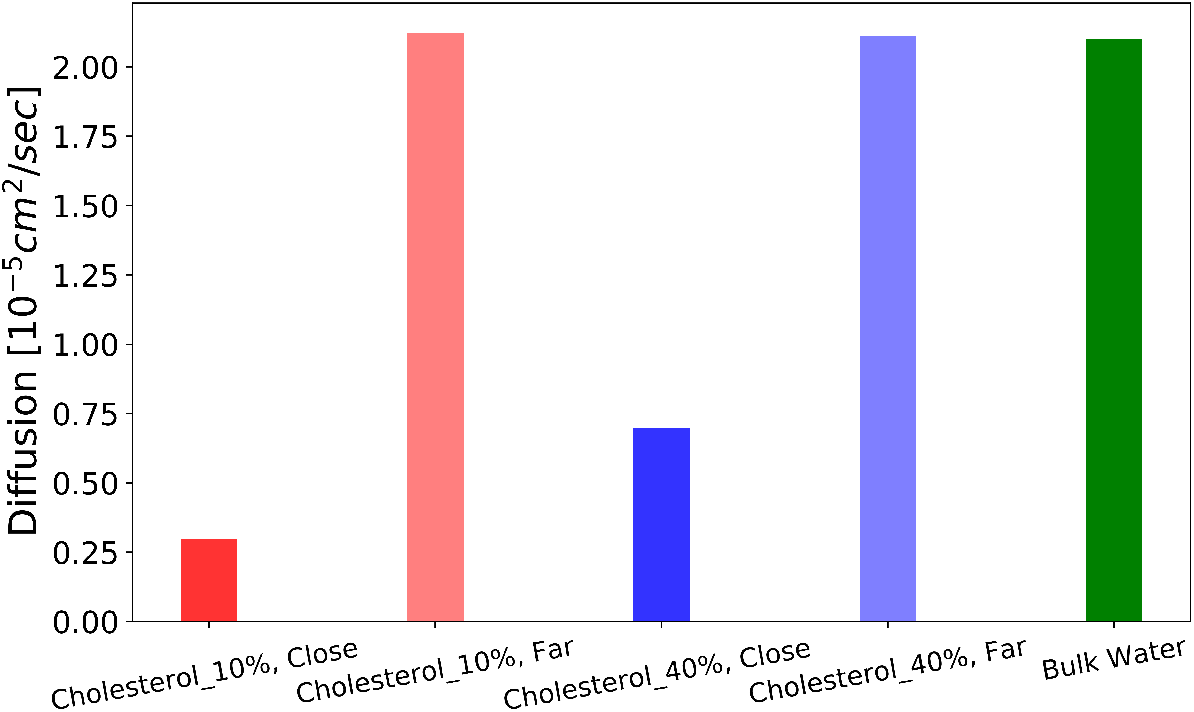
The diffusion coefficient for water molecules close and far from the membranes with different percentages of Cholesterol, in comparison with water diffusion in bulk media

## Conclusion

The results show how cholesterol increases the membrane’s thickness and decreases the surface charge density of the membrane. As a result, water molecules are averagely less attracted to a membrane with a higher cholesterol content. In other words, the number of water molecules interacting with lipid headgroups reduces with the cholesterol content. The dynamical properties, mainly diffusion coefficient, are highly influenced near the membrane due to stronger electrostatic interaction between phospholipids and water molecules. As it is clarified in Fig. 4, the diffusion coefficient of water molecules substantially drops at the vicinity of the membrane relative to far from the surface. This phenomenon is observed more significantly for membranes with less cholesterol content, owing to their stronger electrostatic interactions. A substantial difference in diffusion coefficient near the surface for membranes with various cholesterol content indicates how electrostatic interactions are affected with the introduction of cholesterol. Far from the membrane, as the interaction is less intensive, diffusivity is similar to the bulk case for various Cholesterol contents. Such electrostatic interactions are expected to be more amplified for charged species, such as adenosine triphosphate existing in the biological sub-domains such as extracellular junctions. Previous work [10] showed how confinement and membrane’s electrostatic interaction influence the distribution of nucleotides within extracellular junctions. In [11] it is demonstrated how non-bonding interactions with the surface influence the transport properties of the solvent molecules, particularly in the domains adjacent to the surface. Moreover, in [12], it is determined how electrostatic interaction from the surface charge of crowders such as membrane-bound proteins influence the overall distribution of species. These examples indicate how electrostatic interactions of the surface regulate mass transport properties. Notably, a more attractive membrane to the substrate increases the substrates’ concentration in the intersection. Hence, cholesterol content in the membrane is expected to significantly change species’ distributions inside the extracellular junctions through geometrical and electrostatic effects on the membrane’s surface.

### Molecular dynamic simulation

The original structure of the DMPC membrane is constructed using charm-GUI/membrane builder. Force field parameters of lipid membrane and cholesterol are based on the work of MacKerell et al. [13]. Similar structures are also provided by [14] and a database for different types of lipid membrane. The total number of lipid molecules in each layer of the membrane is 100 so that the percentage indicates the number of cholesterol molecules as well. To create a lipid membrane with a higher number of lipid molecules, like 512 lipid membrane, a 2x2 replication of the 128 lipids in the XY plane with “gromacs” tool “gmx genconf” can be used. The structure was then solvated in an 83A x 83A water box using “packmol,” as shown in Fig. 5. It was minimized and equilibrated after solvation and then was run for 3ns in 293 K in an NVT ensemble. To calculate electron density along the z-axis, we used the density profile tool in VMD [15] which allows us to compute 1-D atomic density along an axis.

**Figure 5:**
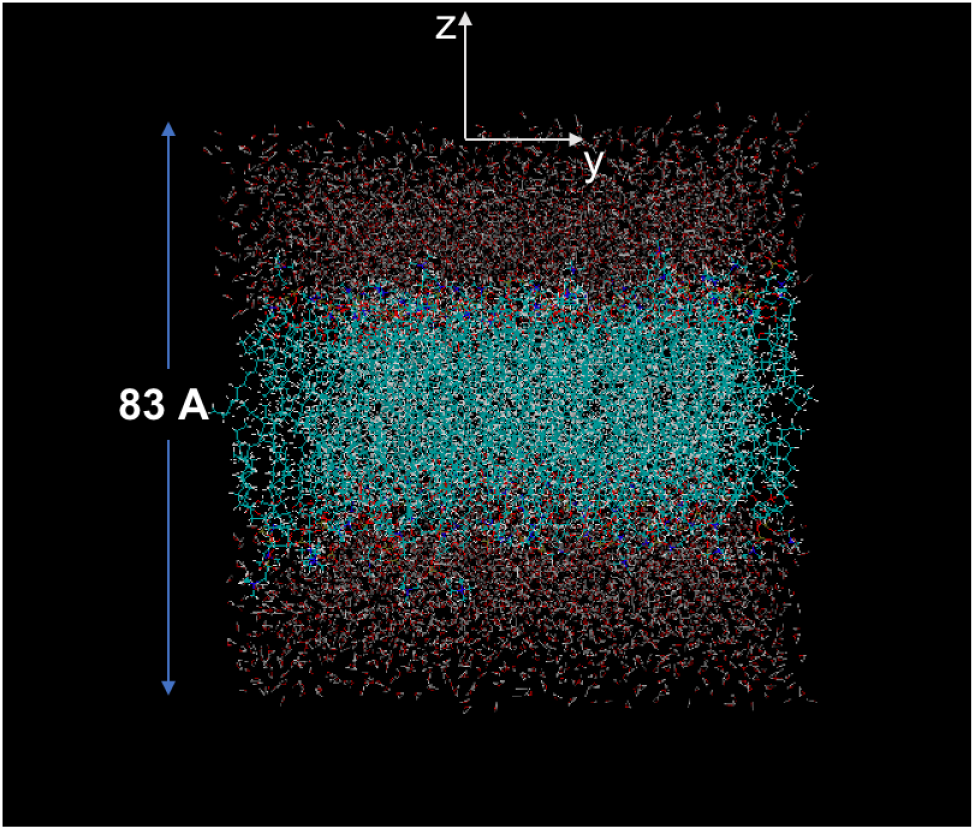
Configuration of the membrane

